# Making headlines: An analysis of US government-funded cancer research mentioned in online media

**DOI:** 10.1101/370973

**Authors:** Lauren A. Maggio, Chelsea L. Ratcliff, Melinda Krakow, Laura Moorhead, Asura Enkhbayar, Juan Pablo Alperin

**Author notes:** Corresponding Post-Publication Author: Juan Pablo Alperin, PhD, Assistant Professor, Simon Fraser University, 515 West Hastings Street, Vancouver, BC, Canada, V6B 5K3. Principle Investigator: Lauren A. Maggio, PhD, MS(LIS), Associate Professor, Department of Medicine, Division of Health Professions Education, Uniformed Services University of the Health Sciences, 4301 Jones Bridge Road, Bethesda, MD 20814.

## Abstract

**Background:** Considerable resources are devoted to producing knowledge about cancer, which in turn is disseminated to policymakers, practitioners, and the public. Online media are a key dissemination channel for cancer research. Yet which cancer research receives media attention is not well understood. Understanding the characteristics of journal articles that receive media attention is crucial to optimize research dissemination.

**Methods:** This cross-sectional study examines journal articles on cancer funded by the US government published in 2016, using data from PubMed and Altmetric to determine whether an article received online media attention. Frequencies and proportions were calculated to describe the cancer types and continuum stages covered in journal articles.

**Results:** 16.8% of articles published on US government-funded research were covered in the media. Published journal articles addressed all common cancers. Roughly one-fourth to one-fifth of journal articles within each cancer category received online media attention. Media mentions were disproportionate to actual burden of each cancer type (ie, incidence and mortality), with breast cancer articles receiving the most media mentions. Cancer prevention and control articles received less online media attention than diagnosis or therapy articles.

**Conclusion:** Findings revealed a mismatch between prevalent cancers and cancers highlighted in the media. Further, journal articles on cancer control and prevention received less media attention than other cancer continuum stages. Media mentions were not proportional to actual public cancer burden nor volume of scientific publications in each cancer category. Results highlight a need for continued research on the role of media, especially online media, in research dissemination.

## INTRODUCTION

The United States (US) federal government is the largest funder of cancer research in the world.[1] Thus, as a public good, it is imperative that the results from federally-funded cancer research be optimally disseminated to all stakeholders, including clinicians, funders, policymakers, and the public. The mass media play a key role in this dissemination.[2-5] To this end, scientific journals have adopted media outreach strategies, such as the *Journal of the National Cancer Institute*’s “Memo to the Media,” to facilitate research dissemination.

Media coverage is critical to health communication across the cancer continuum with particular influence on cancer perceptions and the preventive and screening behaviors among the public.[6-8] Yet little is known about the alignment between the characteristics of federally-funded research articles and how they are covered in the media. This is especially true in light of the expanding nature of the digital media landscape. Such a lack of knowledge impedes the identification and mitigation of any potential discrepancies between the state of the research and what is depicted in the media. Additionally, it disadvantages the public because people may not be exposed to relevant research for making personal health decisions.

In the US, cancer is the most covered disease in the news[9] and the public relies on this coverage as a key source of cancer information.[10, 11] In turn, news reports of cancer research help shape public cancer beliefs,[7, 8, 12] subsequent prevention and detection behaviors,[13, 14] and treatment preferences.[4] Thus, a mismatch between media attention and available evidence can be problematic. For example, past content analyses found breast, blood, and pancreatic cancers to be overrepresented in the news relative to their actual prevalence, while male reproductive, lymphatic, and thyroid cancers were underrepresented relative to prevalence.[5, 15-17] Prevention and detection research also tended to receive less news coverage than other stages of the cancer continuum, such as cancer treatment.[5, 8, 17] While these prior media content analyses provide valuable information, they are based on data more than 15 years old, do not focus specifically on coverage of research and narrowly focus their analysis on selected mainstream news outlets. For example, researchers in 2010 analyzed 13 mainstream print newspapers and magazines from 2005-2007, deciding to exclude online and niche news organizations.[18] Thus, existing research fails to account for today’s broad spectrum of online media that encompasses traditional online news sources as well as trade publications, health and science news aggregators, and press release wire services.

The current study aims to characterize and analyze cancer research articles funded by the US government, including those featured in a collection of more than 2000 online media sources. This analysis provides funders, scientists, and policymakers with an understanding of the dissemination of federally-funded research via online media—information useful in future dissemination and funding initiatives. Additionally, this study provides a replicable method for tracking and analyzing the outputs of funded research across a range of online media, which is critical as traditional news media are no longer the only, nor the primary sources, of health information for the public.[19]

## METHODS

We conducted a cross-sectional study to examine journal articles on cancer funded by the US government and published in 2016. To facilitate transparency and replication of our methods, we have made our computer code and the project’s complete data set publicly accessible at: https://zenodo.org/record/1306985#.W0CYqhJKh24.

We focused on the cancer types most frequently diagnosed in the US, excluding non-melanoma skin cancers. These include cancers of the lung, colon and rectum, pancreas, breast, liver, prostate, leukemia, non-Hodgkin lymphoma, bladder, kidney, endometrium, melanoma, and thyroid (in order of estimated deaths).[20]

To locate articles, we searched PubMed on March 1, 2018, using the Entrez Programming Utilities[21] with a query describing all documents published in journals (including editorials, letters to the editors, etc.) about cancer published in 2016 with any identified source of funding (see Box 1 for search details). We then extracted the record for each document including bibliographic data, funding information, and Medical Subject Heading (MeSH) terms.

**Box 1:**
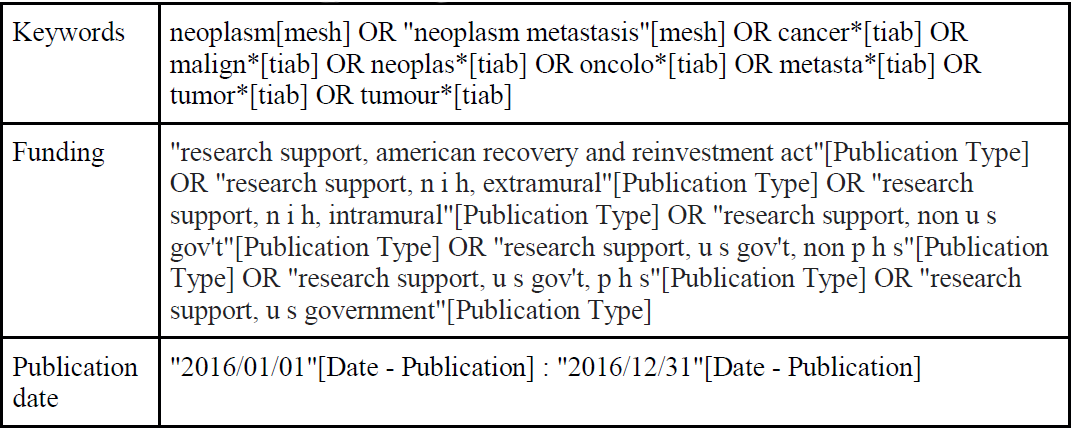
PubMed search strategy for US government funded research on cancer

Articles indexed with MeSH terms for a common cancer or a respective child term were counted as being about that particular cancer (eg, an article indexed with “Leukemia, Plasma Cell” was counted toward the parent term “Leukemia”). When terms for more than one cancer type were included in an article, we counted the article as an occurrence of each cancer. We categorized articles not described by at least one of these prevalent cancer types as “other.” Due to a lack of standardized terminology for stages of the cancer continuum, we partitioned the continuum into stages: prevention and control, diagnosis, and treatment. We utilized corresponding MeSH qualifiers “prevention and control”, “diagnosis”, and “therapy” to identify articles applying to different stages in the cancer control continuum.

After identifying relevant documents, we determined if they had received media attention by searching each document’s PubMed identifier in the database from Altmetric LLC on March 2, 2018. For the purpose of this paper, “media” is defined broadly as online organizations generating and/or aggregating communication. A media “mention” is defined as an instance of either a link to a journal article or a phrase referencing a journal article. Altmetric tracks such mentions of published research, in addition to other impact indicators, such as tweets and Facebook mentions. Altmetric tracks more than 2000 global media sources, including traditional news outlets such as the BBC News, *Los Angeles Times, New York Times*, and *Washington Post*, as well as newer, online-only sources such as *The Daily Beast, Huffington Post,* and *Vox.* Additionally, the company tracks nonprofits and governmental media organizations such as Kaiser Health News, Mayo Clinic, and Voice of America, as well as niche news aggregators and portals such as BioMedReports, EurekAlert, and Health Medicinet. Altmetric also tracks press release news wires such as BusinessWire and PR Newswire. In this study, we consider all these sources in our definition of “media.” (For a listing of media sources, see Altmetric.[22])

### Statistical Analysis

Descriptive statistics were calculated to provide a baseline understanding of the frequency and proportions of journal article characteristics, as well as identify potential differences in the types of cancers and related stages of the cancer continuum covered: prevention/control, diagnosis, and treatment distributed across articles. Subsequently, we examined the frequency and proportions of media mentions received by the articles. Analyses were conducted using SPSS version 21 and Microsoft Excel 365 ProPlus.

## RESULTS

We identified 200264 articles in PubMed on cancer published in 2016. Of these, 11436 (5.7%) reported a US government funding source (all subsequent analysis refers to this subset of n = 11436, unless otherwise specified). The majority of US government-funded articles received funding from the National Institutes of Health (NIH; Table 1) and many received a combination of funding from US government and non-US government sources, such that there were 19944 total funding sources across all articles.

**Table 1:** Number of articles reporting each source of funding support.

| Funding source* | Number of articles |
| --- | --- |
| NIH Extramural | 10377 |
| NIH Intramural | 594 |
| American Recovery & Reinvestment Act | 10 |
| US government Non-Public Health Services | 1922 |
| US government Public Health Services | 295 |
| Non-US government | 6746 |

\*Funding sources defined:
https://www.nlm.nih.gov/pubs/techbull/mj05/mj05_support_heading.html

Articles with multiple funding sources are counted once for each source. All 13 common forms of cancer were the subject of at least one journal article (Table 2). Overall, 41.0% (n = 4687) of journal articles about cancer addressed at least one of these common types of cancer. Most journal articles addressed only one of the top 13 cancers (n = 4448), 195 journal articles addressed 2 top cancers, and 44 journal articles addressed 3-5 top cancers. Frequency of scientific articles differed substantially across the common cancers, with breast cancer (n = 1284), lung cancer (n = 630), and prostate cancer (n = 586) being the subject of the most publications (Table 2; Figure 1).

**Table 2:** Common cancer types covered by journal articles resulting from US government funds in relation to the number of estimated new cases in 2017 and estimated deaths (20).

| Cancer type | Number of scientific articles published in 2016 (n = 4984<sup>†</sup>) | Number of estimated new cases (rank) | Number of estimated deaths (rank) |
| --- | --- | --- | --- |
| Breast | 1284 | 255190* (1) | 41070** (4) |
| Lung | 630 | 222500 (2) | 155870 (1) |
| Colon and Rectal | 535 | 135430 (4) | 50260 (2) |
| Leukemia | 544 | 62130 (9) | 24500 (7) |
| Prostatic | 586 | 161360 (3) | 26730 (6) |
| Pancreatic | 309 | 53670 (12) | 43090 (3) |
| Liver | 302 | 40710 (13) | 28920 (5) |
| Melanoma | 302 | 87110 (5) | 9730 (12) |
| Non-Hodgkin Lymphoma | 170 | 72240 (7) | 20140 (8) |
| Kidney | 106 | 63990 (8) | 14400 (10) |
| Endometrial | 77 | 61380 (10) | 10920 (11) |
| Thyroid | 71 | 56870 (11) | 2010 (13) |
| Urinary/Bladder | 68 | 79030 (6) | 16870 (9) |

<sup>†</sup> Articles may be counted multiple times if they include 2+ cancers
\*Breast cancer total new cases 255190 (252710 female; 2470 male)
\*\*Breast cancer total estimated deaths 41070 (40610 female; 460 male)

**Figure 1:**
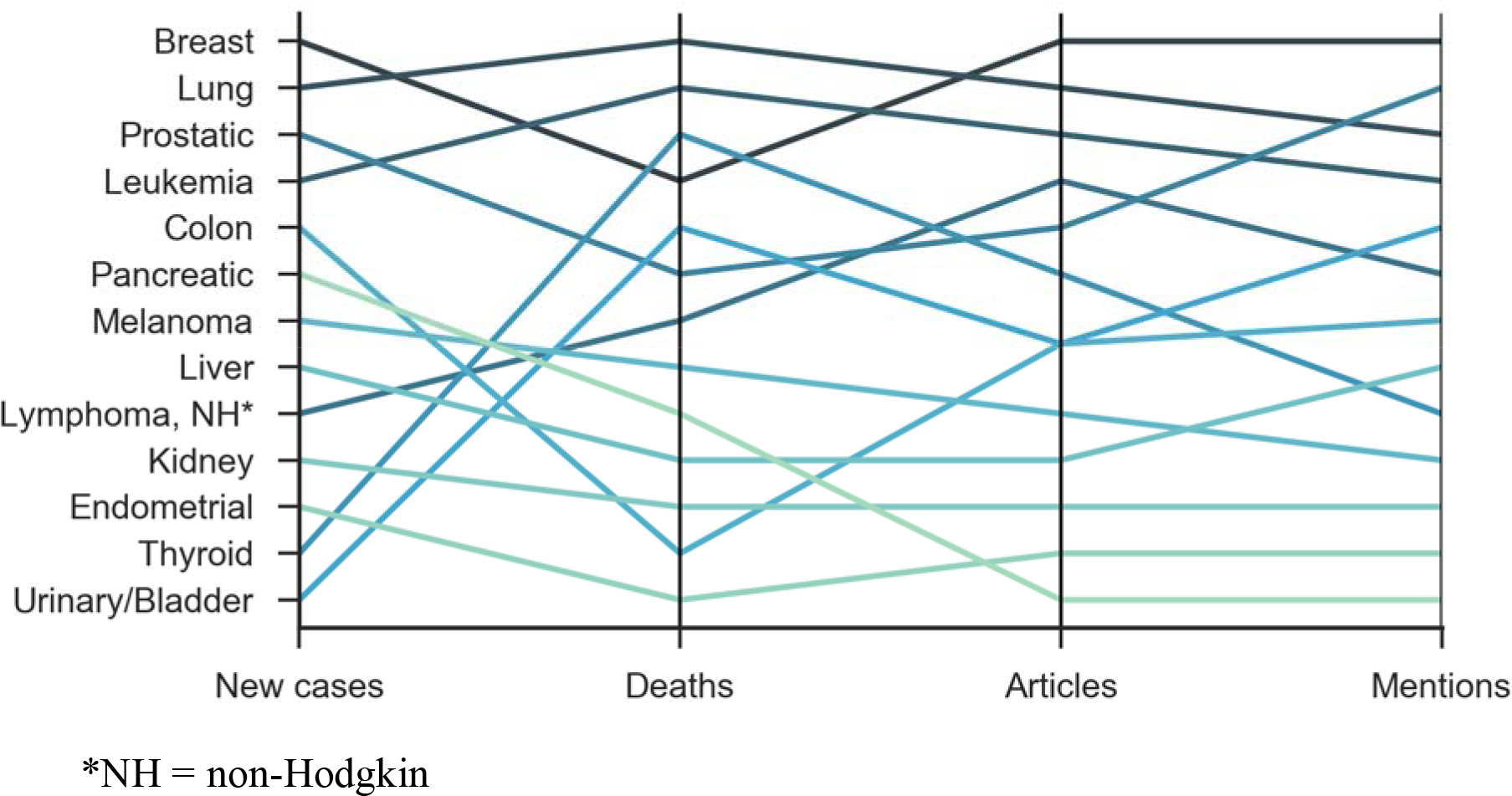
Common cancer types covered by journal articles resulting from US government funds in relation to the number of estimated new cases in 2017 and estimated deaths (20).

US government-funded articles were featured in 1483 unique scientific journals. Articles appeared most frequently in the journals *Cancer Research* (n = 292, 2.6%), *PLOS ONE* (n = 190, 1.7%), and *Proceedings of the National Academy of Sciences (PNAS)* (n = 179, 1.6%). Overall, the 10 journals with the most US government-funded cancer research articles accounted for 14.2% of the total sample of articles (Table 3).

**Table 3:** Top 10 journals featuring the most US government-funded cancer research articles, with and without media coverage.

|  | Journal | Total funded articles in 2016 (% of total articles) | Funded articles with media coverage (% of articles in journal) | Funded articles without media coverage (% of articles in journal) |
| --- | --- | --- | --- | --- |
| 1 | *Cancer Research* | 292 (2.6) | 88 (30.1) | 204 (69.9) |
| 2 | *PloS One* | 190 (1.7) | 20 (10.5) | 170 (89.5) |
| 3 | *Proceedings of the National Academy of the Sciences (PNAS)* | 179 (1.6) | 84 (46.9) | 95 (53.1) |
| 4 | *Oncotarget* | 151 (1.3) | 2 (1.3) | 149 (98.7) |
| 5 | *Blood* | 149 (1.3) | 40 (26.8) | 109 (73.2) |
| 6 | *The Journal of Biological Chemistry* | 147 (1.3) | 10 (6.8) | 137 (93.2) |
| 7 | *Scientific Reports* | 145 (1.3) | 25 (17.2) | 120 (82.8) |
| 8 | *International Journal of Radiation Oncology, Biology, Physics* | 125 (1.1) | 27 (21.6) | 98 (78.4) |
| 9 | *Breast Cancer Research and Treatment* | 118 (1.0) | 10 (8.5) | 108 (91.5) |
| 10 | *International Journal of Cancer* | 112 (1.0) | 13 (11.6) | 99 (88.4) |

Scientific articles also covered the stages of the cancer continuum to varying degrees (Table 4; Figure 2). Across the 13 most common cancer types, 4.4% (n = 206) of articles focused on prevention and control, 11.7% (n = 550) on diagnosis, and 10.7% (n = 502) on therapy.

**Table 4:** Scientific journal articles that identify a stage of cancer continuum.

| Cancer Type | Prevention and control articles (%) | Prevention and control articles with 1+ media mentions (%) | Diagnosis articles | Diagnosis articles with 1+ media mentions (%) | Therapy articles | Therapy articles with 1+ media mentions (%) |
| --- | --- | --- | --- | --- | --- | --- |
| Breast | 55 (4.3%) | 21 (38.2%) | 147 (11.4%) | 30 (20.4%) | 111 (8.6%) | 24 (21.6%) |
| Lung | 25 (4.0%) | 4 (16.0) | 68 (10.8%) | 10 (14.7) | 70 (11.1%) | 12 (17.1) |
| Colon | 53 (9.9%) | 19 (35.8%) | 123 (23.0%) | 27 (22.0) | 45 (8.4%) | 7 (15.6) |
| Leukemia | 18 (3.3%) | 3 (16.7%) | 52 (9.6%) | 11 (21.2) | 87 (16.0%) | 16 (18.4) |
| Liver | 15 (5.0%) | 2 (13.3) | 28 (9.3%) | 1 (3.6) | 35 (11.6%) | 6 (17.1) |
| Prostate | 25 (4.3%) | 8 (32.0) | 68 (11.6%) | 18 (26.5) | 53 (9.0%) | 9 (17.0) |
| Melanoma | 11 (3.6%) | 5 (45.5) | 24 (7.9%) | 6 (25.0) | 44 (14.6%) | 11 (25.0) |
| Pancreas | 11 (3.6%) | 2 (18.2) | 28 (9.1%) | 3 (10.7) | 36 (11.7%) | 4 (11.1) |
| non-Hodgkin<br>Lymphoma | 2 (1.2%) | 0 (0.0) | 21 (12.4%) | 3 (14.3) | 31 (18.2%) | 8 (25.8) |
| Kidney | 1 (0.9%) | 0 (0.0) | 12 (11.3%) | 0 (0.0) | 9 (8.5%) | 2 (22.2) |
| Thyroid | 1 (1.4%) | 0 (0.0) | 7 (9.9%) | 0 (0.0) | 6 (8.5%) | 3 (50.0) |
| Urinary/<br>Bladder | 3 (4.4%) | 1 (33.3) | 12 (17.6%) | 1 (8.3) | 13 (19.1%) | 0 (0.0) |
| Endometrial | 3 (3.9%) | 1 (33.3) | 5 (6.5%) | 0 (0.0) | 8 (10.4%) | (0.0) |
| **Total 13**<br>**common**<br>**cancer types**<br>**(4687)\*** | **206 (4.4%)** |  | **550 (11.7%)** |  | **502**<br>**(10.7%)** |  |
| **Total**<br>**(11436)\*** | **519 (4.5%)** |  | **1000 (8.7%)** |  | **1198**<br>**(10.5%)** |  |

\* Articles may address more than one stage in the cancer continuum. Therefore, these two totals do not reflect unique articles

**Figure 2.**
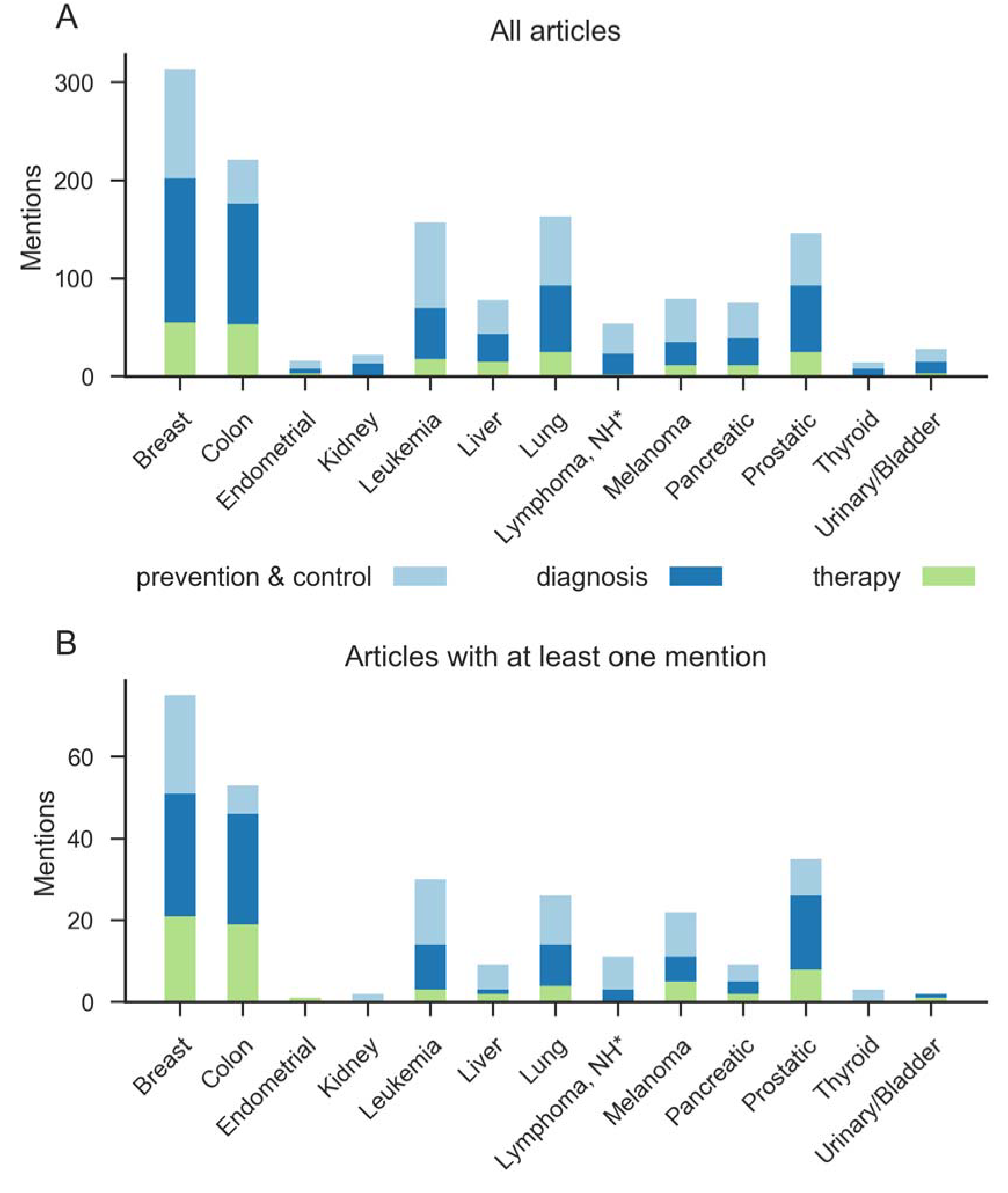
Articles indexed as addressing the prevention and control, diagnosis, or therapy stages of the cancer continuum.

### Mentions in Online Media

A total of 1925 (16.8%) of cancer research articles were mentioned in online media at least once. The majority (n = 9511, 83.2%) of journal articles did not receive a media mention (median = 0, mean = 1.84, SD = 11.98, range = 0-462). Of the 1925 articles that did receive at least one media mention: 735 received just one mention (6.4%), 198 received 2 mentions (1.7%), and 992 (8.8%) received 3 or more mentions. Four articles received over 300 mentions (Table 5). [23-32]

**Table 5:** Top 10 articles by media mentions

| Rank | Journal | Article title | # media mentions |
| --- | --- | --- | --- |
| 1. | *Lancet* | Does physical activity attenuate, or even eliminate, the detrimental association of sitting time with mortality? A harmonised meta-analysis of data from more than 1 million men and women[23] | 462 |
| 2. | *Oncogene\** | Targeting MET and AXL overcomes resistance to sunitinib therapy in renal cell carcinoma[24] | 355 |
| 3. | *JAMA Dermatology* | The risk of cancer in patients with psoriasis: A population-based cohort study in the Health Improvement Network[25] | 346 |
| 4. | *JAMA* | Screening for Colorectal Cancer: US Preventive Services Task Force Recommendation Statement[26] | 317 |
| 5. | *Science* | Mutational signatures associated with tobacco smoking in human cancer[27] | 244 |
| 6. | *Nature* | Naturally occurring p16(Ink4a)-positive cells shorten healthy lifespan[28] | 221 |
| 7. | *Journal of Psychopharmacology* | Rapid and sustained symptom reduction following psilocybin treatment for anxiety and depression in patients with life-threatening cancer: a randomized controlled trial[29] | 200 |
| 8. | *JAMA* | CDC Guideline for Prescribing Opioids for Chronic Pain — United States, 2016[30] | 183 |
| 9. | *Science* | The phenotypic legacy of admixture between modern humans and Neanderthals[31] | 178 |
| 10. | *JAMA Internal Medicine* | Association of religious service attendance with mortality among women[32] | 176 |

\**Oncogene* is a Nature Journal

The frequency of media mentions differed across cancer types (Table 6; Figure 3). The proportion of journal articles, by cancer type, receiving at least one media mention ranged from 7.0% of thyroid cancer articles to 24.8% of melanoma articles. Thus, while breast cancer had the largest *total number* of articles with one or more media mentions (251), melanoma articles received the largest *relative proportion* of coverage (24.8%). In other words, while breast cancer research may have the widest reach in terms of absolute number of online media mentions, a larger proportion of journal articles on melanoma are being mentioned online.

**Table 6:**
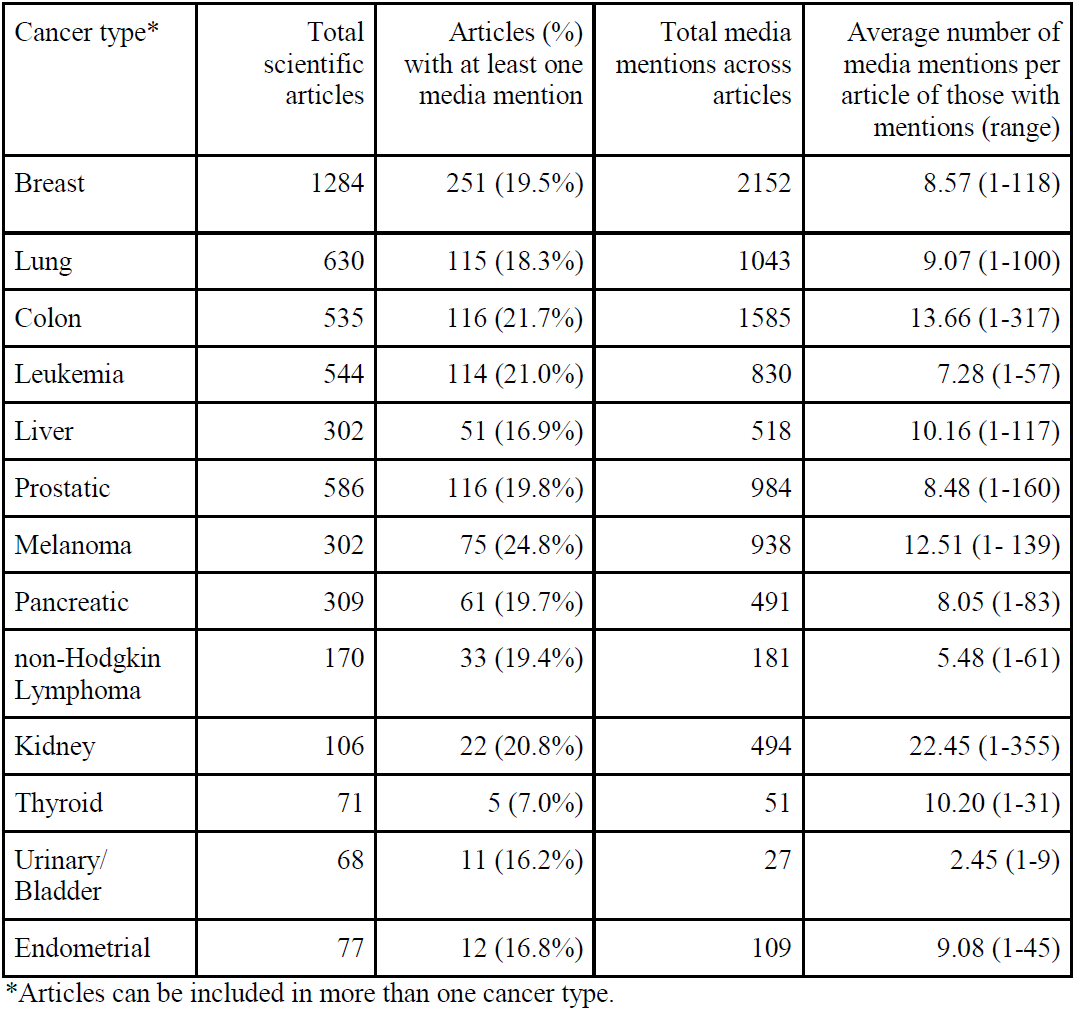
Number of scientific articles about common cancer types in relation to media mentions

| Cancer type* | Total scientific articles | Articles (%) with at least one media mention | Total media mentions across articles | Average number of media mentions per article of those with mentions (range) |
| --- | --- | --- | --- | --- |
| Breast | 1284 | 251 (19.5%) | 2152 | 8.57 (1-118) |
| Lung | 630 | 115 (18.3%) | 1043 | 9.07 (1-100) |
| Colon | 535 | 116 (21.7%) | 1585 | 13.66 (1-317) |
| Leukemia | 544 | 114 (21.0%) | 830 | 7.28 (1-57) |
| Liver | 302 | 51 (16.9%) | 518 | 10.16 (1-117) |
| Prostatic | 586 | 116 (19.8%) | 984 | 8.48 (1-160) |
| Melanoma | 302 | 75 (24.8%) | 938 | 12.51 (1- 139) |
| Pancreatic | 309 | 61 (19.7%) | 491 | 8.05 (1-83) |
| non-Hodgkin Lymphoma | 170 | 33 (19.4%) | 181 | 5.48 (1-61) |
| Kidney | 106 | 22 (20.8%) | 494 | 22.45 (1-355) |
| Thyroid | 71 | 5 (7.0%) | 51 | 10.20 (1-31) |
| Urinary/ Bladder | 68 | 11 (16.2%) | 27 | 2.45 (1-9) |
| Endometrial | 77 | 12 (16.8%) | 109 | 9.08 (1-45) |

\*Articles can be included in more than one cancer type.

**Figure 3:**
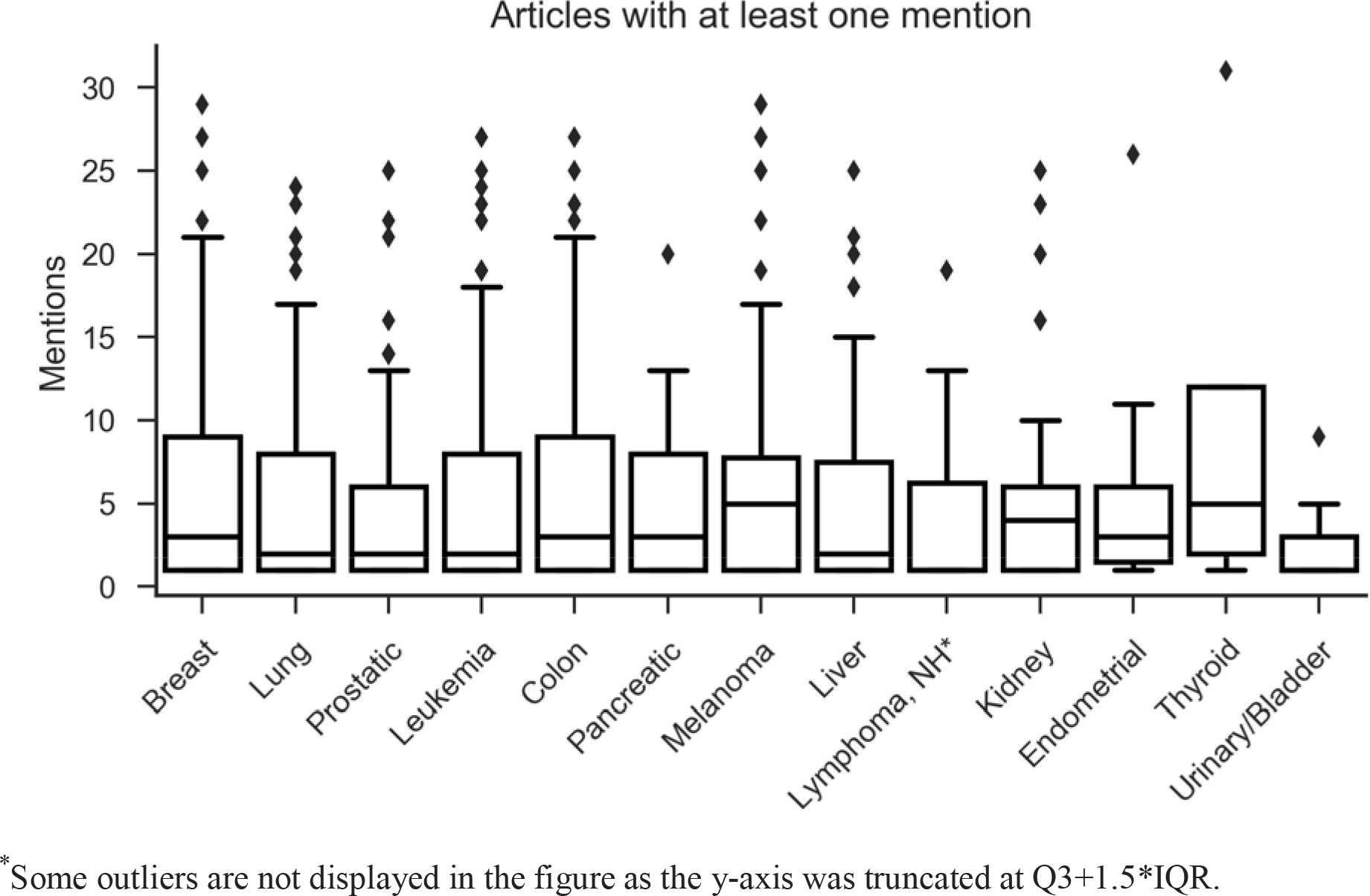
Articles with at least one news mention by cancer type.*

Media coverage also varied across cancer continuum stage for the top 13 cancers. While prevention and control articles received more relative coverage (32.0%), compared to diagnosis (20.0%) or treatment (20.3%), this stage had fewer absolute mentions (66 mentions) compared to other stages (110 and 102 mentions, respectively), translating into lower absolute dissemination for the earliest stage of the continuum.

## DISCUSSION

Media coverage is a key access point to cancer information for the public.10, 11] Approximately one out of every six articles reporting a US government funding source received media attention. For the public to receive useful information to guide health, it is imperative that the information disseminated to the public be accurately interpreted and aligned with available evidence.[33] Our findings provide stakeholders a valuable formative landscape of the current dissemination of federally-funded cancer research captured across a spectrum of online media attention, including traditional news sources as well as trade publications, health and science news aggregators, and press release wire services. Looking to the future, our approach provides funder groups, such as the NCI and NIH, with a replicable method for tracking outputs from their research portfolio as they appear in online media.

### Alignment with previous literature

While journal articles spanned all common cancer types, they were not represented in proportions mirroring estimates of cancer burden. Similar to earlier findings,[5] breast cancer was the most-published-on cancer type, with more than double the frequency of the second-most focused-on cancer, despite breast cancer causing fewer deaths than lung, colon and rectal, and pancreatic cancer. Prior research found a similar mismatch between the cancers prominent in the scientific literature and those with the highest actual burden (ie, prevalence, incidence, or mortality).[5, 15-17] This misalignment is noteworthy and the public, physicians, policymakers, and the media should understand that the research volume does not necessarily indicate the severity or population-level burden of the disease. It should also be noted that a complex set of factors, including investigator-initiated research proposals and available funding, influences scientific research.

Published articles disproportionately represented the stages of the cancer continuum, with prevention and control research accounting for a smaller proportion than diagnosis or treatment research. This echoes prior research findings[5, 8, 17] highlighting an underrepresentation of prevention-focused research. Notably, there were a number of cancers (eg, lung, melanoma) with a surprisingly low amount of prevention/control research represented in this sample.

Media mentions were somewhat comparable across the 13 common cancers: Around one-fifth to one-quarter of journal articles on each cancer type received media attention (with the exception of thyroid cancer, which was substantially lower). However, similar to publication volume, the volume of media mentions per cancer type was disproportionate to actual incidence of each cancer type and varied across the cancer continuum. Breast cancer was the most mentioned cancer type in online media, mirroring the higher volume of published scientific articles about breast cancer. This aligns with prior research that found certain cancers, including breast cancer, were overrepresented in news coverage relative to incidence and mortality rates.[5] Although fewer scientific articles on melanoma were published during this same period, melanoma research had the highest amount of *relative* media attention, or percent of articles receiving media mentions (24.8%). We also found limited media attention to cancer prevention and detection research across the top 13 cancer types, consistent with past studies.[5, 15]

### Limitations

Due to the nature of our methods, we may have inadvertently excluded articles. For example, we included only articles indexed in MEDLINE, which omits articles published in journals not indexed there. Additionally, authors or publishers may have failed to report funding sources; however, the NIH reports 89% compliance with the requirement to deposit manuscripts.[34] In relation to content, we recognize that the US government also funds basic research that while in the future may greatly influence clinical cancer research, at this time would not be specifically indexed as cancer-related and therefore not retrieved. It is also possible that some articles related to cancer prevention/control (eg, topics such as tobacco cessation interventions) did not include relevant MeSH (eg, neoplasm) constituting our search criteria. Additionally, articles containing MeSH for more than one cancer type were included in counts across multiple categories, which may affect estimation. Media mention data were collected by Altmetric, which searches online media for mentions of articles based on unique article IDs (eg, PubMed IDs), links to publisher websites, and a series of proprietary techniques. Altmetric’s approach, to our knowledge, offers the most comprehensive tracking of research in online media, but it remains imperfect, with some errors readily identifiable and an unknown number of missed mentions. Furthermore, while Altmetric searches over 2000 sources, other potentially important sources may not be included.

More broadly, while online media outlets are important channels for disseminating health information, we recognize there are other impactful channels, such as social media campaigns, that warrant study.[3] Further, while we focused on the characteristics of scientific journal articles receiving media mentions, it is also important to consider how such research is covered and portrayed in the news media—notably online, where readership shows signs of growth— suggesting a future research opportunity.

### Implications

Our findings provide a unique understanding of the dissemination of federally-funded cancer research in online media via an extensive collection of media sources. This benchmark can be used to evaluate future dissemination and funding initiatives that take into consideration the new information landscape that reaches beyond traditional news media. Second, these results can inform scientists as to the characteristics of cancer research mentioned in the media. Third, these findings provide funder groups, such as the NCI and NIH, with a replicable method for future tracking of the outputs from their research portfolio that appear in the online news media.

Media mentions of research can impact the public’s health care utilization, physicians’ practices, and research funding agendas.[35] For example, researchers credit a rapid decline in use of hormone replacement therapy by women and their physicians to the news media coverage of a randomized trial linking the therapy to potential harms.[36, 37] Additionally, media coverage can influence which research focus (eg, cancer types, cancer stages) is likely to receive federal funding, public policy attention, or scholarly/scientific attention. A 2014 analysis found that discrepancies in media coverage and in public estimates of cancer were mirrored in federal funding for cancer research (eg, overrepresentation of breast and blood cancers and underrepresentation of bladder cancer, relative to actual incidence.[7]) This highlights an important area for future examination; data and methods from this study could facilitate such efforts.

Researchers have found that online media attention to scientific articles about cancer treatment correlates positively with the presence of media outreach.[38, 39] Thus, unsurprisingly the top 10 articles receiving the most media attention were published in journals that engage in outreach and dissemination activities. For example, four of the articles appear in the JAMA Network. The JAMA Network promises prospective authors a dedicated media team and formal press office that provides journalists early access to articles, writes press releases, and creates video and audio recordings of scientists discussing their findings. Similar services are provided by the Lancet, Science, and Nature publishing groups, which account for all but one of the top ten articles. This suggests an important role for targeted media outreach in ensuring that a broader range of scientific research receives online media coverage. Scientists, universities, and smaller scientific journals may wish to model these media outreach practices. Additionally, a need likely exists for more affordable outreach and dissemination services.

Media are recognized as an important channel for knowledge dissemination.[3] However, online media generally have received little attention as a communication channel in dissemination models and frameworks. Future research should examine how online media could be optimally incorporated into dissemination processes and knowledge translation strategies. Specifically, it will be useful to understand how marketing/communications teams can work with researchers to connect media more directly with emerging scientific research.

As a strength of this research, we included a broad mix of online media organizations, including traditional news media, broadcast organizations, trade publications, health and science news aggregators, and press release wire services and public relations platforms. This broad inclusion is critical as traditional news media are no longer the only sources, nor the primary sources, of health information for the public.[19]

Thus, our findings provide a unique understanding of dissemination of the results of federally-funded cancer research in the broader landscape of online media. This benchmark can be used to evaluate future dissemination and funding initiatives that take into consideration the new information landscape that reaches far beyond traditional news media.

## Disclaimer

The views expressed in this article are those of the authors and do not necessarily reflect the official policy or position of the Uniformed Services University of the Health Sciences, the Department of Defense, the National Cancer Institute, or the US Government.

